# Different biological effects of exposure to far-UVC (222 nm) and near-UVC (254 nm) irradiation

**DOI:** 10.1101/2022.10.28.514223

**Authors:** Renata Spagolla Napoleão Tavares, Douglas Adamoski, Alessandra Girasole, Ellen Nogueira Lima, Amauri da Silva Justo-Junior, Romênia Domingues, Ana Clara Caznok Silveira, Rafael Elias Marques, Murilo de Carvalho, Andre Luis Berteli Ambrosio, Adriana Franco Paes Leme, Sandra Martha Gomes Dias

## Abstract

Ultraviolet C (UVC) light has long been used as a sterilizing agent, primarily through devices that emit at 254 nm. Depending on the dose and duration of exposure, UV 254 nm can cause erythema and photokeratitis and potentially cause skin cancer since it directly modifies nitrogenated nucleic acid bases. Filtered KrCl excimer lamps (emitting mainly at 222 nm) have emerged as safer germicidal tools and have even been proposed as devices to sterilize surgical wounds. All the studies that showed the safety of 222 nm analyzed cell number and viability, erythema generation, epidermal thickening, the formation of genetic lesions such as cyclobutane pyrimidine dimers (CPDs) and pyrimidine-(6-4)-pyrimidone photoproducts (6-4PPs) and cancer-inducing potential. Although nucleic acids can absorb and be modified by both UV 254 nm and UV 222 nm equally, compared to UV 254 nm, UV 222 nm is more intensely absorbed by proteins (especially aromatic side chains), causing photooxidation and cross-linking. Here, in addition to analyzing DNA lesion formation, for the first time, we evaluated changes in the proteome and cellular pathways, reactive oxygen species formation, and metalloproteinase (MMP) levels and activity in full-thickness *in vitro* reconstructed human skin (RHS) exposed to UV 222 nm. We also performed the longest (40 days) *in vivo* study of UV 222 nm exposure in the HRS/J mouse model at the occupational threshold limit value (TLV) for indirect exposure (25 mJ/cm^2^) and evaluated overall skin morphology, cellular pathological alterations, CPD and 6-4PP formation and MMP-9 activity. Our study showed that processes related to reactive oxygen species and inflammatory responses were more altered by UV 254 nm than by UV 222 nm. Our chronic *in vivo* exposure assay using the TLV confirmed that UV 222 nm causes minor damage to the skin. However, alterations in pathways related to skin regeneration raise concerns about direct exposure to UV 222 nm.

## 1 Introduction

Ultraviolet (UV) light is electromagnetic radiation comprising three wavelength bands: UV-A (400-315 nm), UV-B (315-280 nm), and UV-C (280-100 nm) [1]. In addition to vacuum UV-C (100-200 nm), far-UV-C incorporates wavelengths between 200 and 230 nm [2], while near UV-C incorporates wavelengths between 230 and 280 nm. The well-known germicidal effect of UV-C light is based on nucleic acid mutagenesis, which decreases viability, reduces clonogenic survival, and induces apoptosis and cytogenetic damage [1]. This effect can inactivate bacteria, bacterial spores, fungi, viruses, and protists on object surfaces, in water, and in air **[1,3–5]**.

In this context, mercury vapor-based UV lamps emitting predominantly at 254 nm are used in UV germicidal irradiation (UVGI) systems. UVGI systems can be found in heating and ventilation devices and air conditioning equipment and are often used in hospitals and other public spaces **[6–9]**. Nevertheless, UV light causes premutagenic UV-associated DNA lesions, such as cyclobutane pyrimidine dimers (CPDs), and pyrimidine-(6-4)-pyrimidone photoproducts (6-4PPs) in the skin and eyes and induces oxidative stress via the generation of reactive oxygen species (ROS) [10]. In addition, human exposure to near UVGI (> 230 nm) systems has been proven to be harmful to corneal cells of the eyes and the outermost layer of skin cells, leading to transitory photokeratitis and erythema, respectively [11].

More recently, far-UV-C lamps (< 230 nm) have emerged as safe tools to sanitize populated environments, as the light they generate is recognized to be absorbed by the skin stratum corneum with little effect on the genetic material of living cells [12]. Recent advances in the development and production of filtered KrCl excimer lamps with shorter peak wavelengths (222 nm) hold promise in this regard [12,13]. However, according to the physical properties of UV-C, UV light at 222 nm interacts with many carbon bonds and aromatic rings, potentially crosslinking aromatic amino acids and, for that reason, has the potential to promote alterations in the proteome of mammalian cells, with the potential to affect cell and tissue function [14].

To define safe limits of UV exposure, the American Conference of Governmental Industrial Hygienists (ACGIH) established a threshold limit value (TLV), defined as the dose to which a worker can be indirectly exposed eight hours a day, 40 hours per week without adverse health effects [11]. The International Commission on Non-Ionizing Radiation Protection (ICNIRP), in collaboration with the World Health Organization (WHO), has adopted these same guidelines [11]. As UV radiation ranges from 100 to 400 nm, the TLV varies according to wavelength. The TLV for UV 254 nm is 6.0 mJ/cm², while for UV 222 nm, the dose limit is 25.0 mJ/cm² for indirect human exposure.

Safety studies of KrCl excimer lamps have mainly evaluated the capacity of 222 nm light to induce DNA damage in eukaryotic cells using *in vitro* and *in vivo* models [12,13,15–17]. Several studies concluded that human and animal cells could tolerate far-UV-C with little DNA dimer formation [18], which was shown to disappear after 24 h of exposure [19], did not induce erythema even at the highest human tested dose [16,20], and did not increase cell death or inflammatory responses mediated by cytokines **[12,16,19,21]**. Recently, one study used electron paramagnetic resonance (EPR) spectroscopy to show that irradiation with 40 mJ/cm^2^ UV 222 nm can generate radical species on irradiated reconstructed human skin (RHS) **[19]**.

To date, no study has performed a more in-depth evaluation of the molecular alterations that occur in skin tissues irradiated with UV 222 nm, especially regarding protein alterations. Additionally, no previous study considered occupational TLV exposure to thoroughly understand the effects of 222 nm on mammalian skin, especially after several days of exposure. To better understand the molecular effects of UV 222 nm on the skin, we used the RHS model, which recapitulates the physiology of full-thickness human skin (dermis and epidermis); we performed proteomics analysis to evaluate altered cellular pathways and detected and quantified reactive oxygen species formation, as well as the level and activity of metalloproteinases. We have also performed a 40-day/8-hour per day exposure of the hairless HRS/J mouse model to either UV 254 nm or UV 222 nm at their respective TLV doses. We showed that UV 222 nm causes fewer molecular, cellular, and tissue alterations than UV 254 nm.

## 2 Materials and Methods

### 2.1 Reconstructed human skin model (RHS)

Normal human dermal fibroblasts (CRL-2703 #lot2584882) were obtained from ATCC and cultivated in Iscove’s modified Dulbecco’s medium (IMDM) (ATCC® 30-2005™) supplemented with 10% fetal bovine serum (FBS, Invitrocell, Brazil). Normal human keratinocytes (nh-skp-KT0037) were obtained from *Banco de Células do Rio de Janeiro* and cultivated in Dermal Cell Basal Medium (ATCC® PCS-200-030™) supplemented with a Keratinocyte Growth Kit (ATCC® PCS-200-040™). Both types of skin cells originated from the foreskin of boys younger than 10 years after medically indicated circumcision (CAAE: 46338521.3.0000.0123). The full-thickness (epidermis and dermis) RHS was developed in-house in a 24-well plate size insert (Millicell-PCF, Merck, Brazil), and it was fully differentiated after ten days, as described previously **[22,23]**.

### 2.2 HRS/J hairless model

Six-week-old female hairless mice (HRS/J; Campo Grande, Brazil) were used in this study. Mice were maintained under specific pathogen-free conditions at the LNBio Bioterium, CNPEM. All experiments were carried out in accordance with the Guidelines for Animal Experimentation of CNPEM. The experimental protocol was approved by the ethics committee of the LNBio, CNPEM (Protocol: 91).

### 2.3 UV-C devices and irradiation assays

Two test apparatuses were designed, optimized, fabricated, and calibrated to enable accurate and controlled UV-C treatment (at 254 nm or 222 nm) [24] of the RHS samples and acute irradiation of the mice. The RHS was kept within the inserts and placed in the middle of a 30 mm Petri dish containing 3 mL of Dermal Cell Basal Medium (ATCC® PCS-200-030™). The irradiance levels of the lamps inside the treatment chamber were measured using calibrated UV-C sensor systems, namely, an HD2302.0 Delta OHM system with an LP 471 UV-C (260 nm) detector (Instruterm) or a UIT2400 Handheld Light Meter 222 nm radiometer (Ushio), which provided irradiance patterns and levels from which optimal treatment locations and time could be determined. The RHS was snap frozen immediately (0 h), 24 h or 48 h post-irradiation. The RHS collected 24 h or 48 h post-irradiation was placed, along with its insert, in a 24-well plate containing 0.5 mL of Dulbecco’s modified Eagle’s medium – DMEM high glucose (Vitrocell, Brazil) supplemented with 10% fetal bovine serum (FBS) (Vitrocell, Brazil) and incubated at 37 °C and 5% CO_2_. For acute irradiation of mice, we anesthetized the animals with an intraperitoneal injection of ketamine (100 mg/kg) and xylazine (10 mg/kg), and the animals were placed inside the apparatus at a predefined position. The animals were euthanized by cervical dislocation, and a 5 cm² sample of dorsal skin was removed with scissors and tweezers.

We built a device for the *in vivo* chronic irradiation assay, which was placed inside the animal cage. The device was 3D printed by extrusion using polylactic acid filament in an S3X Printer (Sethi3D) to create four compartments, within which one animal was placed during the irradiation period. Each mouse had access to water and food through specifically printed dispensers. The lamp was placed at the top of the device, within a 3D-printed box, controlled by an electronic household timer to limit exposure to the light. The irradiation area of the lamp was equally distributed among the four compartments due to the compartment-specific mesh pattern added to the front of the lamp, guaranteeing that each animal received similar UV doses. Each mouse was exposed to a total of 3.125 mJ/cm²/hour of UV 222 nm or 0.75 mJ/cm²/hour of UV 254 nm for eight hours daily. Control animals (n=4) were kept inside a similar device without a lamp. The animals were irradiated five days a week at random positions with two days off, for a total of 40 days of irradiation. The total time of the assay was 56 days. We measured the animals’ weight daily. The dorsal skin were collected following the last day of the experiment.

### 2.4 Tissue analysis by light microscopy

The RHS was snap-frozen in OCT Compound (Tissue-Tek, Sakura). The frozen material was sectioned into 8 µm slices using a cryostat (LEICA CM1950, Germany). After hematoxylin and eosin staining of cryosections, immunofluorescence (IF) or immunohistochemistry (IHQ) was performed. The dorsal skin of the mice was removed and fixed with 4% phosphate-buffered methanol-free formaldehyde for 24 hours. After 20 min under tap water, the specimens were dehydrated in an ethanol gradient ranging from 70% to 100% in 10% increments, with the following combinations: ethanol:xylene, xylene, Paraplast Plus (Sigma):xylene, and pure Paraplast Plus. Tissue sections with a thickness of 8 μm were obtained using a microtome (LEICA RM2255, Germany). After deparaffinization and rehydration of the mouse skin tissue sections, we conducted hematoxylin and eosin (HE) staining. HE staining was conducted as reported previously [25], with Harris hematoxylin (Laborclin, Brazil) and 2% eosin solution (InLab, Brazil). Before IF or IHQ of mouse dorsal skin slices, we performed an antigen retrieval process by incubating the sections with citrate buffer inside a receptacle with ice and warmed them with five cycles of 1 min of microwave heating at maximum potency.

To characterize the RHS, we performed IF using rabbit antibodies against keratin-10 (1:200, ab76318, Abcam) or keratin-14 (1:200, ab7800, Abcam). First, we circled the sections with a hydrophobic pen and fixed them with 4% methanol-free formaldehyde in phosphate buffered saline (PBS – 137 mM NaCl, 2.7 mM KCl, 10 mM Na_2_HPO_4_, 1.8 mM KH_2_PO_4_) for 10 min, rinsed the sections with PBS, and blocked them for one hour with 10% goat serum in PBS. Next, we aspirated the liquid and applied the primary antibodies for one hour at room temperature; we rinsed with PBS and applied a secondary goat anti-rabbit Alexa 546 or 488 (Invitrogen) antibody for one hour at room temperature and protected from light, then rinsed and counterstained with 4′,6-diamidino-2-phenylindole (DAPI) (D9542, Sigma□Aldrich). The same IF protocol was applied to the other antibodies. IF of metalloproteinases and metallopeptidase inhibitor 1 was performed with primary anti-MMP1 (1:100; ab137332), anti-TIMP1 (1:250; ab216432) and anti-MMP9 [RM1020] (1:100; ab283575) antibodies (Abcam, EUA). Secondary goat anti-rabbit Alexa Fluor 546 or goat anti-rabbit Alexa Fluor 488 antibodies (1:1000; Invitrogen) were incubated for one hour, and then the samples were rinsed and counterstained with DAPI blue (D9542, Sigma□Aldrich).

To detect DNA lesions, IF and IHQ were performed with anti-cyclobutane pyrimidine dimer (CPD) (1:500 IF, 1:200 IHC, Cosmo, Clone TDM-2, CAC-NM-DND-001) or anti-6-4-pirimidine-pirimidone photoproduct (6-4-PP) (1:500 IF; 1:200 IHC, Cosmo, Clone 64M-2, CAC-NM-DND-002) antibodies for one hour. After incubation with primary antibody, the slides were rinsed with PBS and incubated with goat anti-mouse Alexa Fluor 546 or goat anti-mouse Alexa Fluor 488 (Invitrogen) for one hour.

To perform IHC, the sections were blocked with 5% goat serum and 5% BSA in PBS for one hour, and afterward, the sections were blocked with 5% hydrogen peroxide for 5 min. We incubated the slides with secondary anti-mouse HRP-conjugated (GE, Healthcare) overnight at 4 °C. After rinsing with PBS, the color reaction was developed by adding diaminobenzidine (ImmPACT DAB Peroxidase HRP Substrate, Vector Biolabs). Following this, counterstaining was performed with Harris hematoxylin (Laborclin). Images were captured with a fluorescence microscope (DM6, Leica, Germany) using the HC PL FLUOTAR 10x/0.32 and HC PL FLUOTAR 20x/0.55 lenses and analyzed with LasX software (Leica, Germany). CPD- and 6,4-PP-positive cells were quantified using a script developed in ImageJ/Fiji (10.1038/nmeth.2019) and R software, available on GitHub (https://github.com/douglasadamoski/farUVC).

### 2.5 Mass spectrometry analysis

The epidermis of the RHS was mechanically removed, rinsed with 1 mL PBS, and solubilized with 50 µL of 100 mM Tris-HCl pH 7.4, 8 M urea, 2 M thiourea, and 1 µM EDTA. We cleaned-up the samples by the addition of ice-cold acetone (8 volumes) and methanol (1 volume), followed by the incubation of samples overnight at −20 °C. After that, we added 5 mM DL-dithiothreitol (DTT, Sigma□Aldrich) for 25 min at 56 °C, followed by the addition of 14 mM iodoacetamide (IAA, Sigma Aldrich) for 30 min at room temperature protected from light [25]. After these steps, we added 1 mM calcium chloride (Sigma□Aldrich), followed by digestion with 0.2 µg trypsin (Sequencing Grade Modified Trypsin, V5111, Promega) for 16 hours at 37 °C. After digestion with trypsin, the reaction was stopped by adding 1% formic acid (Merck), pH < 3. Then, the samples were desalted using Stage Tips with C18 membranes (Octadecyl C18-bonded silica - 3M Empore extraction disks) and thoroughly dried in an evaporator (SPD 1010 SpeedVac, Thermo) **[26]**. One aliquot containing 800 ng of Bradford-quantified peptides was analyzed on an ETD-enabled Orbitrap Velos mass spectrometer (Thermo Fisher Scientific, Waltham, MA, USA) connected to the EASY-nLC system (Proxeon Biosystem, West Palm Beach, FL, USA) through a Proxeon nanoelectrospray ion source. Peptides were separated by a 2–90% acetonitrile gradient in 0.1% formic acid using PicoFrit analytical column (20 cm x ID75 μm, 5 μm particle size, New Objective) at a flow rate of 300 nL·min−1 over 212 min. The nanoelectrospray voltage was set to 2.2 kV, and the source temperature was 275 °C. All instrument methods were used in the data-dependent acquisition mode. The full-scan MS spectra (m/z 300–1600) were acquired in the Orbitrap analyzer after accumulation to a target value of 1 × 10^6^. The resolution in the Orbitrap was set to r = 60,000, and the 20 most intense peptide ions with charge states ≥2 were sequentially isolated to a target value of 5,000 and fragmented in the linear ion trap using low-energy CID (normalized collision energy of 35%). The signal threshold for triggering an MS/MS event was 1,000 counts. Dynamic exclusion was enabled with an exclusion size list of 500, an exclusion duration of 60 s, and a repeat count of 1. An activation q = 0.25 and activation time of 10 ms was used (Kawahara et al., 2014). The identification of proteins was performed with MaxQuant v.1.5.8 [27] using the UniProt Human Protein Database (release January 2022; 100,731 sequences; 40,968,421 residues). Carbamidomethylation was set as a fixed modification, and N-terminal acetylation and methionine oxidation were set as variable modifications, with a maximum of 1 missed trypsin cleavage, and tolerance values of 10 ppm for precursor mass and 1 Da for fragment ions were set for protein identification. We used a filter to achieve a maximum false discovery rate of 1% at the peptide and protein levels, using a reverse target-decoy database strategy with reverse peptide sequences as decoy entries **[26]**.

We filtered proteins reverse- and only-identified-by-site, and single replicate entries. Data were log_2_-transformed, and missing not at random (MNAR) values were defined intra-treatment as proteins with a mean below the 5% percentile and replaced using the minimum replacement method [28]. All other missing values were defined as missing completely at random (MCAR) and further processed by Bayesian principal component analysis value imputation (bPCA, [29]. Data were normalized by the upper-quantile method [30] and used as input for differential abundance analysis using limma [31–33]. Pathways from the Molecular Signatures Database (MSigDB, [34] were used as input to calculate the single-sample score, as implemented in singScore [35]. When a molecular signature had two elements, designated “_DN” and “_UP,” scores were calculated considering those expression direction differences.

### 2.6 ROS assay

We incubated RHS with 50 µM DCFH_2_-DA diluted in PBS for 45 min at 37 °C in 5% CO_2_. After washing with PBS, the tissues were irradiated, while control tissues were kept in the dark. Immediately after the irradiation period, the total fluorescence intensity of each tissue was measured in an EnSpire microplate reader (Perkin-Elmer) adapted to a 24-well size plate and then snap-frozen in OCT compound (TissueTek, Sakura) for sectioning. Histological sections with a thickness of 8 µm were obtained with a cryostat (LEICA CM1950) and analyzed with a Leica TCS SP8 confocal mounted on a Leica DMI 6000 inverted microscope. Images were processed using LAS AF Lite software (Leica, Germany), and fluorescence signals were quantified using Python scripts **[23,36]**.

### 2.7 Zymography

Mouse skin (5 mm^2^) was lysed with 500 µL of 50 mM Tris-HCl at pH 7.4, 150 mM NaCl, 2 mM EGTA, 1 µM pepstatin, 10 mM sodium orthovanadate, and 1% Triton x-100 solution for 5 min with the aid of a tissue homogenizer (Polytron PT2100, Kinematica). The sample was centrifuged at 12,000 x g and 4 °C, and the supernatant was collected and quantified with a Bradford (Bio-Rad) reaction kit. Twenty micrograms of each sample was subjected to electrophoresis through prepared 10% SDS-polyacrylamide gels containing 0.1% acidic digestion of collagen (porcine gelatin, Sigma, G1890) as a substrate [37] at 150 mV for 2 h. Next, the gel was washed with water and incubated for one hour with 50 mM Tris-HCl, 5 mM CaCl_2,_ 1 µM ZnCl_2, and_ 2.5% Triton x-100 solution to achieve renaturation of the proteins. After this period, the gel was incubated overnight at 37 °C with 50 mM Tris-HCl, 5 mM CaCl_2_, and 1 µM ZnCl_2_ to allow metalloproteinase activity. Each gel was then stained with 0.5% Coomassie blue and destained with water.

## 3 Results

### 3.1 UV 222 nm produces less DNA damage than UV 254 nm *in vitro* and *in vivo*

We used the previously described in-house device to perform the studies with the *in vitro* RHS model and HRS/J mice (short-term exposure) [21]. The RHS model is an organotypic three-dimensional *in vitro* culture composed of primary human skin cells (fibroblasts and keratinocytes). The RHS recapitulates the physiology of full-thickness human skin, namely, the dermis and epidermis. In the supra basal layer, the expected stratification of the epidermis through the cornified layer mimics the barrier function of the skin **[38,39]**. After eight days of differentiation of the RHS at the air-liquid interface, the models were morphologically characterized by immunofluorescence of cytokeratin 10 (supra basal layer) and 14 (basal layer) and HE staining (Supplementary Fig 1). As expected, the RHS was fully differentiated, presenting a proliferative basal layer, stratum spinosum, granulosum, and corneum (Supplementary Fig 1).

The RHS was nonirradiated or irradiated with 500 and 1500 mJ/cm² doses of UV 222 nm or UV 254 nm, and the material was immediately frozen (0 h) or incubated for an additional 24 h or 48 h (herein called acute exposure) to assess changes in morphology, desquamation/regeneration of the cell layers, and the formation of CPD and 6-4-PP. As an immediate effect of irradiation (0 h), we verified that only a 1500 mJ/cm² dose of UV 254 nm could induce desquamation of the RHS (Figure 1a). Immunohistochemistry and immunofluorescence with anti-CPD and anti-6,4-PP revealed that UV 222 nm induced the generation of less CPD and 6,4-PP than UV 254 nm at the same doses at 0 h (Figure 1a and d), as previously reported [16,20,21]. Interestingly, at 24 h (Figure 1b and d) or 48 h (Figure 1c and d) post-irradiation, the genetic lesions were much more persistent in the RHS irradiated with UV 254 nm than in that irradiated with UV 222 nm. At 48 h, while the RHS exposed to UV 254 nm irradiation presented pronounced desquamation of the stratum corneum and the granulosa layer, the RHS exposed to UV 222 nm lamp presented desquamation only in the stratum corneum, with apparent epidermal thickening, likely related to a regenerative process (Figure 1c).

**Fig. 1.**
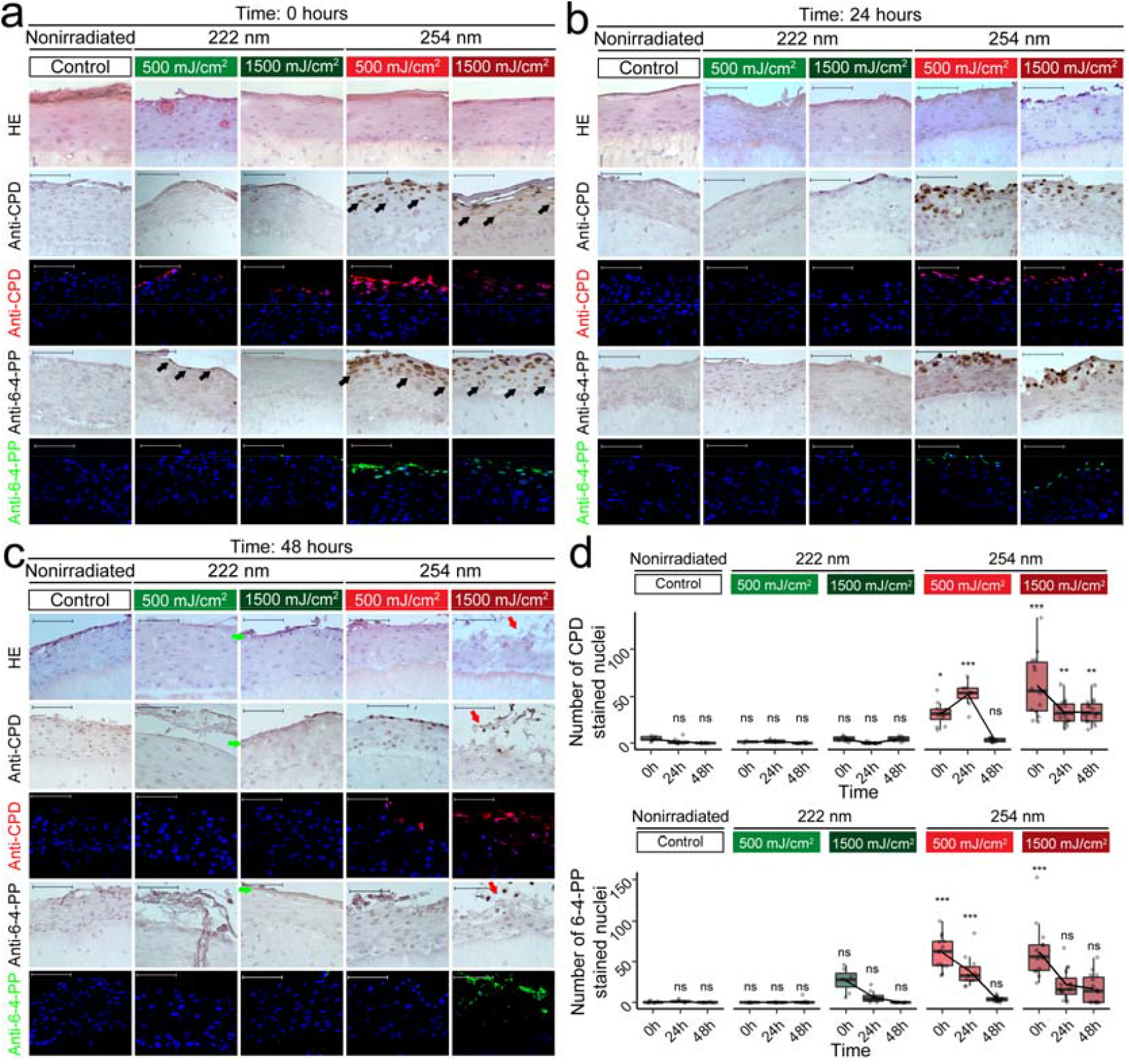
UV 222 nm produces fewer DNA lesions than UV 254 nm in the RHS. The RHS was nonirradiated or irradiated with 500 or 1500 mJ/cm2 UV 222 nm or 254 nm and evaluated at 0 h (a), 24 h (b) or 48 h (c) post-exposure. HE: hematoxylin and eosin staining. Anti-CPD and anti-6-4-PP staining were evaluated by immunohistochemistry (brownish coloration) or immunofluorescence (anti-CPD in red and anti-6,4-PP in green; nuclei were stained with 4’,6-diamidino-2-phenylindole DAPI, in blue). (d) Immunohistochemistry quantified nuclei positive for anti-CPD or anti-6-4-PP were quantified in the nonirradiated RHS, or the RHS irradiated with 500 or 1500 mJ/cm^2^ UV 222 nm or 254 nm at 0 h, 24 h or 48 h post-irradiation. Positive nuclei for immunohistochemistry staining are indicated by black arrows. Skin desquamation by 1500 mJ/cm^2^ dose of 254 nm at 48 h is indicated by red arrows; green arrows indicate skin thickening caused by UV 222 nm exposure 48 h post-exposure. Images are representative of the treatment group. P-values were derived from ANOVA followed by Tukey’s honestly significant difference test comparing all conditions to timepoint 0 h from nonirradiated control. *p < 0.05, **p < 0.01, and ***p < 0.0001.

We then investigated the effect of 40 days of exposure for eight hours a day (hereafter called chronic exposure) to TLV doses of UV 222 nm and UV 254 nm, namely, 25 and 6 mJ/cm², respectively, in terms of the formation of DNA lesions in the skin of HRS/J mice. To perform the assay, we built a device that separated the animals into four compartments, with one animal per space, and evaluated the irradiation throughout the space they occupied (Supplementary Fig 2a-f). The animals received a total dose of 1000 mJ/cm² UV 222 nm or 240 mJ/cm² UV 254 nm. As a control for the experiment, we acutely exposed anesthetized mice to 6 mJ/cm² (UV 254 nm), 25 mJ/cm² (UV 222 nm), or 1500 mJ/cm² (UV 222 and 254 nm), euthanized them and collected the skin. As expected, we observed anti-CPD- and anti-6,4-PP-positive nuclei in the skin of the mice irradiated with 1500 mJ/cm² UV 254 nm but not UV 222 nm (Supplementary Fig 3). Surprisingly, even 6 mJ/cm² UV 254 nm produced a detectable dimer signal, while 25 mJ/cm² UV 222 nm did not produce any detectable staining using either immunofluorescence or immunohistochemistry techniques (Supplementary Fig 3). After 40 days of exposure to either 6 mJ/cm² (UV 254 nm) or 25 mJ/cm² (UV 222 nm), no sunburn or desquamation was observed in the dorsal skin of mice irradiated with either UV-C lamp (Supplementary Fig 4a). To further investigate the effect of chronic irradiation with UV 222 and 254 nm on the epidermis of mouse skin, histological analyses were performed (Supplementary Fig 4b). We did not observe parakeratosis, epidermal hyperplasia, intracellular edema, or mitotic bodies in the stratum spinosum of skin irradiated with UV 222 nm (data not shown)[40]. Histological analysis showed no changes in the architecture of the skin irradiated 40 days with UV 222 nm and UV 254 nm groups compared with nonirradiated group. We did not observe thickening of the epidermis, evidence of spongiosis, or hyperkeratosis [40]. Dermal compartment did not show inflammatory foci or increased thickening of the extracellular matrix [40]. Nonetheless, we observed hydropic degeneration focus characterized by nucleus enlargement, chromatin condensation, evident nucleoli, and reduced cytoplasm in addition to areas of epidermal atrophy in the acute 222 nm group (Supplementary Fig 4b) [40]. The hydropic degeneration described was intensified in the acute 254 nm group compared to the acute 222 nm group (Supplementary Fig 4b). Importantly, we could not detect any CPD or 6,4-PP signal in the irradiated mouse skin (Figure 2). In conclusion, UV 222 nm induced fewer DNA lesions than UV 254 nm, and these lesions were more superficial and readily eliminated by desquamation.

**Fig. 2.**
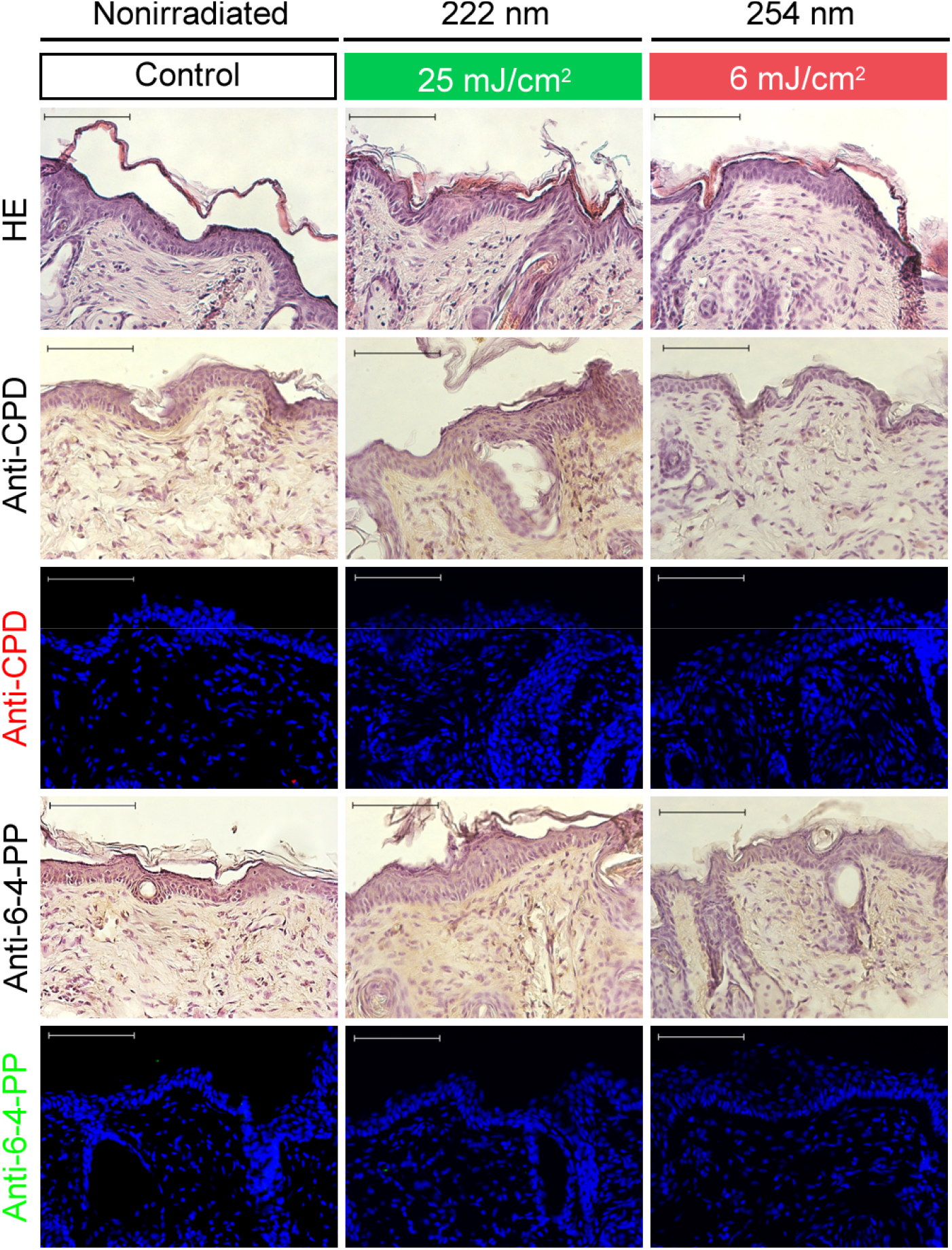
Chronic exposure of HRS/J mice to the TLVs of UV 222 nm and 254 nm did not cause morphological changes or DNA lesions in the skin. Mice were nonirradiated or irradiated with either 6 mJ/cm^2^ UV 254 nm or 25 mJ/cm^2^ UV 222 nm for eight hours a day, five days a week, for a total of 40 days. HE: hematoxylin and eosin staining. Anti-CPD and anti-6-4-PP staining were evaluated by immunohistochemistry (brownish coloration) or immunofluorescence (anti-CPD in red and anti-6,4-PP in green; nuclei were stained with DAPI in blue). Images are representative of the treatment group.

### 3.2 Proteomics analysis of the RHS model showed that irradiation with UV 222 nm leads to less alteration of extracellular matrix, oxidative stress, and inflammatory response pathways than UV 254 nm

Since acute 1500 mJ/cm² or chronic exposure to TLV of UV 222 nm doses (cumulative dose of 1000 mJ/cm²), respectively, induced few or no detectable DNA lesions, we asked about the alterations in protein levels (and cellular pathways) induced by UV 222 nm in the skin epidermal layer, especially in comparison to UV 254 nm. The RHS model was nonirradiated or irradiated with 1500 mJ/cm² UV at 222 nm or 254 nm, and then the samples were analyzed at 0 h, 24 h, or 48 h post-exposure. The epidermis was detached, and the proteome was analyzed by mass spectrometry. After removing potentially misidentified and contaminating peptides (Supplementary Fig 5a), total LFQ counts were not significantly different among the processed samples (Kruskal□Wallis p-value 0.85, Supplementary Fig 5b). A combination method was employed to deal with missing peptides not at random (MNAR), using minimum value replacement [28], and missing peptides completely at random (MCAR), using Bayesian PCA imputation [29], followed by log-transformation of upper-quantile normalized quantification values. As expected, this approach yielded comparable protein abundances among replicates and experimental conditions (Supplementary Fig 5c-d).

Since skin morphological changes were more visible at 48 h post-irradiation, we used this treatment group as the focus of our analysis. We detected 193 differentially abundant proteins (p-value <0.05, limma) between 1500 mJ/cm² exposure to 222 nm and 254 nm after 48 h of irradiation (Figure 3a, Supplementary Table). Interestingly, unsupervised cluster analysis grouped the control samples (nonirradiated) closer to the samples irradiated with UV 222 nm and more distant from the samples irradiated with UV 254 nm, showing that UV 254 nm induced more changes in the epidermis than UV 222 nm (Figure 3a, Supplementary Table).

**Fig. 3.**
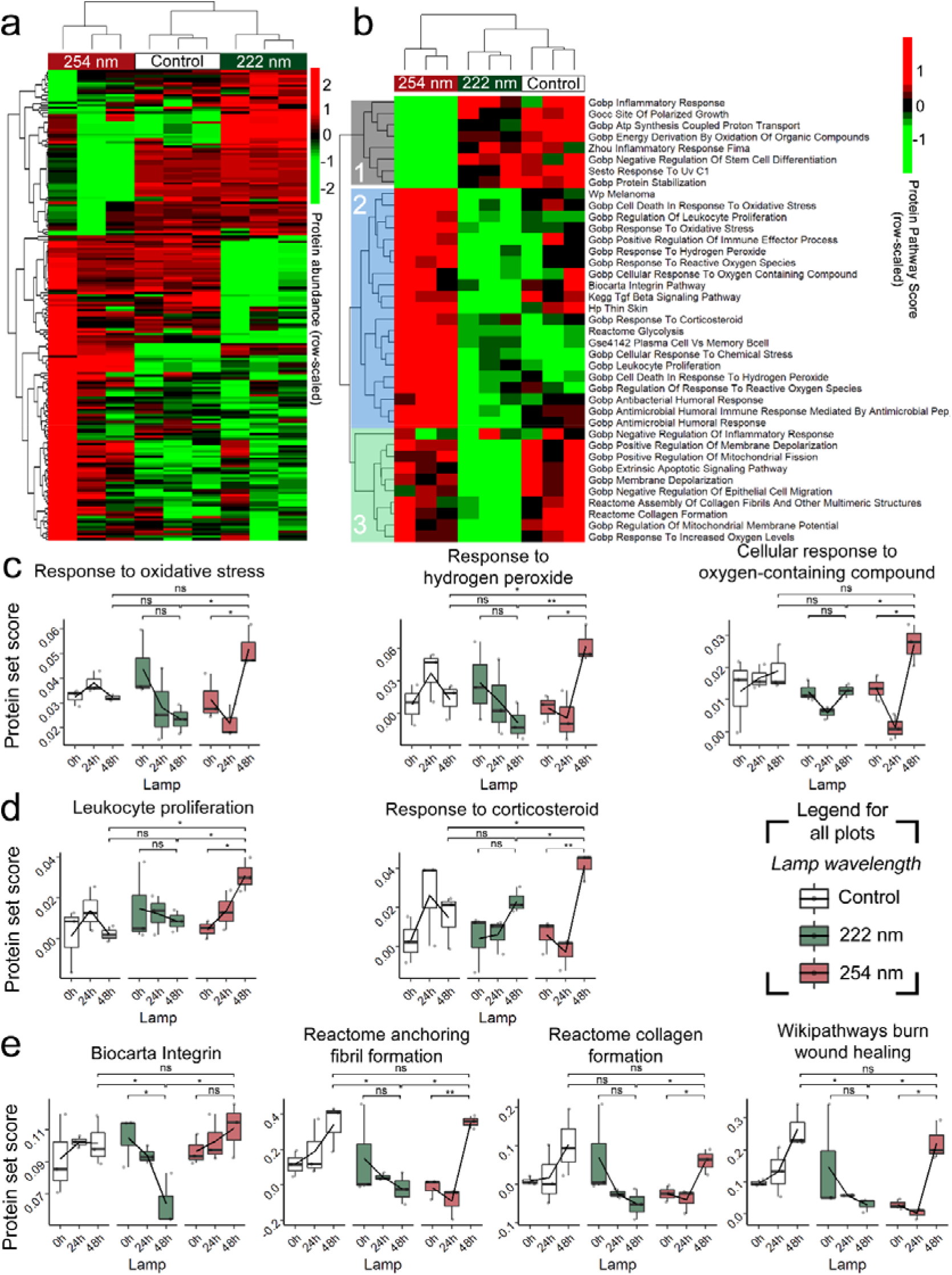
Proteomics analysis of the RHS shows that both UV 222 nm and 254 nm promote protein pathway alterations. (a) Heatmap of 193 differentially abundant proteins (p-value <0.05, limma) between 1500 mJ/cm² exposure to UV 222 nm and UV 254 nm after 48 h of irradiation. Values are row-scaled imputed log2 upper-quantile normalized label-free quantifications from MaxQuant. (b) Heatmap of selected pathways involved in oxidative stress, inflammation, and the extracellular matrix. The values are protein pathway scores defined by the sample. For the grouping scheme, see the main text. For complete heatmap, see Supplementary Figure 6. For both heatmaps, the unsupervised clustering method was complete, and the distance was defined by Pearson’s correlation. (c) Time-resolved protein set scores for gene ontology biological processes pathways related to “response to oxidative stress”, “response to hydrogen peroxide”, and “cellular response to oxygen-containing compounds”. (d) Time-resolved protein set scores for gene ontology pathways related to “leukocyte proliferation” and “response to corticosteroids”. (e) Time-resolved protein set scores for “integrin pathways” (Biocarta), “anchoring fibril formation” (Reactome), “collagen formation” (Reactome), and “burn wound healing” (Wikipathways). For (c-e), the upper and lower limits of the boxplot represent 0.75 and 0.25 quantile estimations, respectively, whiskers extend to the limits of the values, and the dark horizontal line denotes the median. Lines connect the means of each time point. Each dot represents an individual cell or sample. P-values were derived from Welch’s two-sided t-test. *p < 0.05, **p < 0.01, and ***p < 0.0001.

MSigDB analysis revealed 635 pathways with varying protein scores (p-value < 0.05, t-test) between UV 254 nm and 222 nm 48 h post-irradiation (Supplementary Fig 6A and Supplementary Table 2). Several of these pathways were related to skin and extracellular matrix organization, oxidative stress, and inflammatory response (Supplementary Table 2). The clustering of these pathways reproduced the separation seen for the differentially abundant proteins across the treatments (Figure 3b). Moreover, clustering according to pathway scores distinguished three groups: Group 1, in which the scores were similar between the control and UV 222 nm but lower for UV 254 nm; Group 2, in which the scores were similar between the control and UV 222 nm but higher for UV 254 nm; and Group 3, in which the scores were similar between the control and UV 254 nm but lower for UV 222 nm.

A time-course analysis of the pathways found in Group 2 revealed, for UV 222 nm treatment, unaltered “response to oxidative stress”, “response to hydrogen peroxide” and “cellular responses to oxygen-containing compounds” scores over time (Figure 3c), in contrast to UV 254 nm, which induced the highest scores for all three pathways at 48 h post-exposure (Figure 3c). Additionally, in Group 2, regarding general inflammatory pathways, skin exposed to UV 222 nm did not present variations in the “leucocyte proliferation” or “response to corticosteroid” scores, while UV 254 nm induced the highest scores 48 h after exposure (Figure 3d). Importantly, in Group 3, “Biocarta integrin”, “Reactome anchoring fibril formation,” “Reactome collagen formation”, and “Wikipathways burn wound healing” showed a pattern in which scores decreased after 48 h of exposure to UV 222 nm (only significant for Biocarta integrin) but increased for the UV 254 nm treatment (non significant for Biocarta integrin), implying that these processes may be suppressed after irradiation with UV 222 nm (Figure 3e). Taken together, these results suggest that UV 222 nm generated less oxidative stress and a reduced inflammatory response but could disturb processes related to skin regeneration.

### 3.3 Irradiation with UV 222 nm generated a smaller increase in skin metalloproteinase levels than UV 254 nm

Since the proteomic analysis revealed that UV 222 nm has the potential to affect skin regeneration, we evaluated the levels and activity of metalloproteinases in skin exposed to this type of radiation. Matrix metalloproteinases (MMPs) are zinc-containing endopeptidases with a wide range of substrate specificities. Collectively, these enzymes can degrade various components of the extracellular matrix (ECM). MMP-1 is a collagenase, while MMP-9 is a gelatinase [41,42]. Tissue inhibitor of metalloproteinase-1 (TIMP-1) inhibits most metalloproteases, including MMP-1 and MMP-9 [41,42]. The alterations to the ECM induced by MMPs might contribute to skin wrinkling, a characteristic of premature skin aging. Compared to those in the nonirradiated control, MMP-1 and MMP-9 levels were increased in the dermis of RHS irradiated with 500 or 1500 mJ/cm² doses of both UV 222 nm and UV 254 nm after 48 h of exposure; qualitatively, the signal intensity seemed higher in the dermis and epidermis of the RHS irradiated with UV 254 nm than in the RHS irradiated with UV 222 nm (Figure 4). Interestingly, while TIMP-1 was detected in the dermis of the RHS irradiated under all tested conditions, the signal intensity seemed higher in the RHS (both dermis and epidermis) irradiated with either a 500 or 1500 mJ/cm² dose of UV 222 nm, suggesting more pronounced activation of skin defense and repair processes compared to that observed after irradiation with UV 254 nm (Figure 4).

**Fig. 4.**
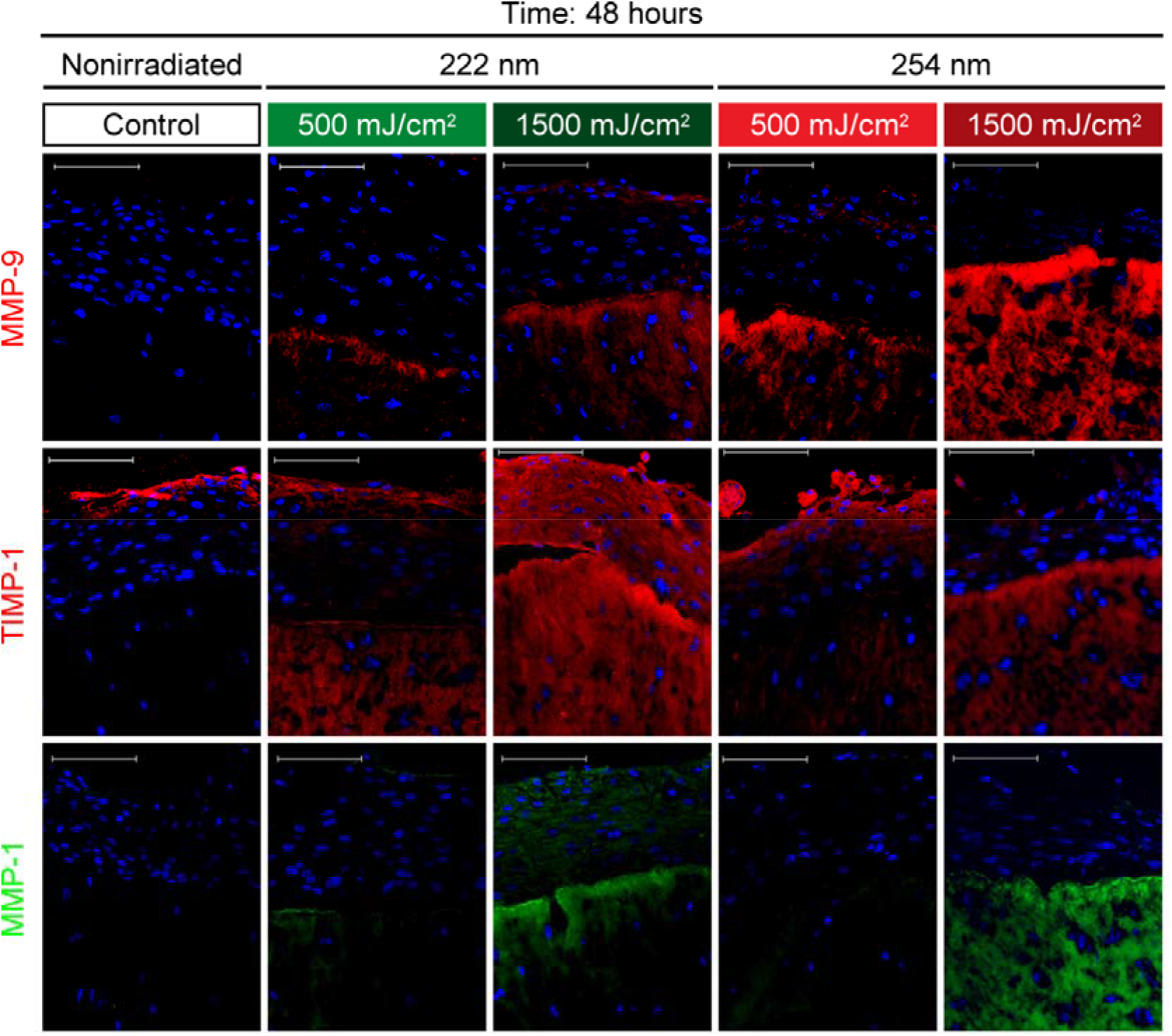
Acute UV 222 nm and UV 254 nm irradiation led to increased levels of MMP-1, MMP-9, and TIMP-1 in the RHS. The RHS was nonirradiated or irradiated with 500 or 1500 mJ/cm^2^ UV 222 nm or UV 254 nm and was evaluated at 48 h post-exposure. Anti-MMP-1, anti-MMP-9, and anti-TIMP-1 staining were evaluated by immunofluorescence (anti-MMP-1 in green and anti-MMP-9 and anti-TIMP-1 in red; nuclei were stained with DAPI (in blue)).

Finally, we assessed the effect of chronic irradiation of mouse skin with 6 mJ/cm² UV 254 nm or 25 mJ/cm² UV 222 nm on MMP-1, MMP-9, and TIMP-1 levels. Under these experimental conditions, at the end of 40 days of exposure, only MMP-9 levels were increased in the epidermis of animals exposed to either of these types of UV irradiation (Figure 5a). We also used a zymography assay to identify the active form of MMP-9 after chronic irradiation of the mice. While all four mice exposed to 6 mJ/cm² UV 254 nm presented strong precursor (pre) and active forms of MMP-9 bands at the end of the assay, two of the mice irradiated with 25 mJ/cm^2^ of UV 222 nm and three of the nonirradiated control mice presented a weaker active form of MMP-9 bands (Figure 5b). In conclusion, acute exposure to 1500 mJ/cm² UV 222 nm can increase metalloproteinase levels in the skin. However, chronic exposure to the TLV dose of UV 222 nm (1000 mJ/cm² total dose) was less effective than UV 254 nm in increasing the levels of active MMP-9, indicating that this type of radiation is safer regarding to leading to extracellular matrix degradation at the occupational safety level.

**Fig. 5.**
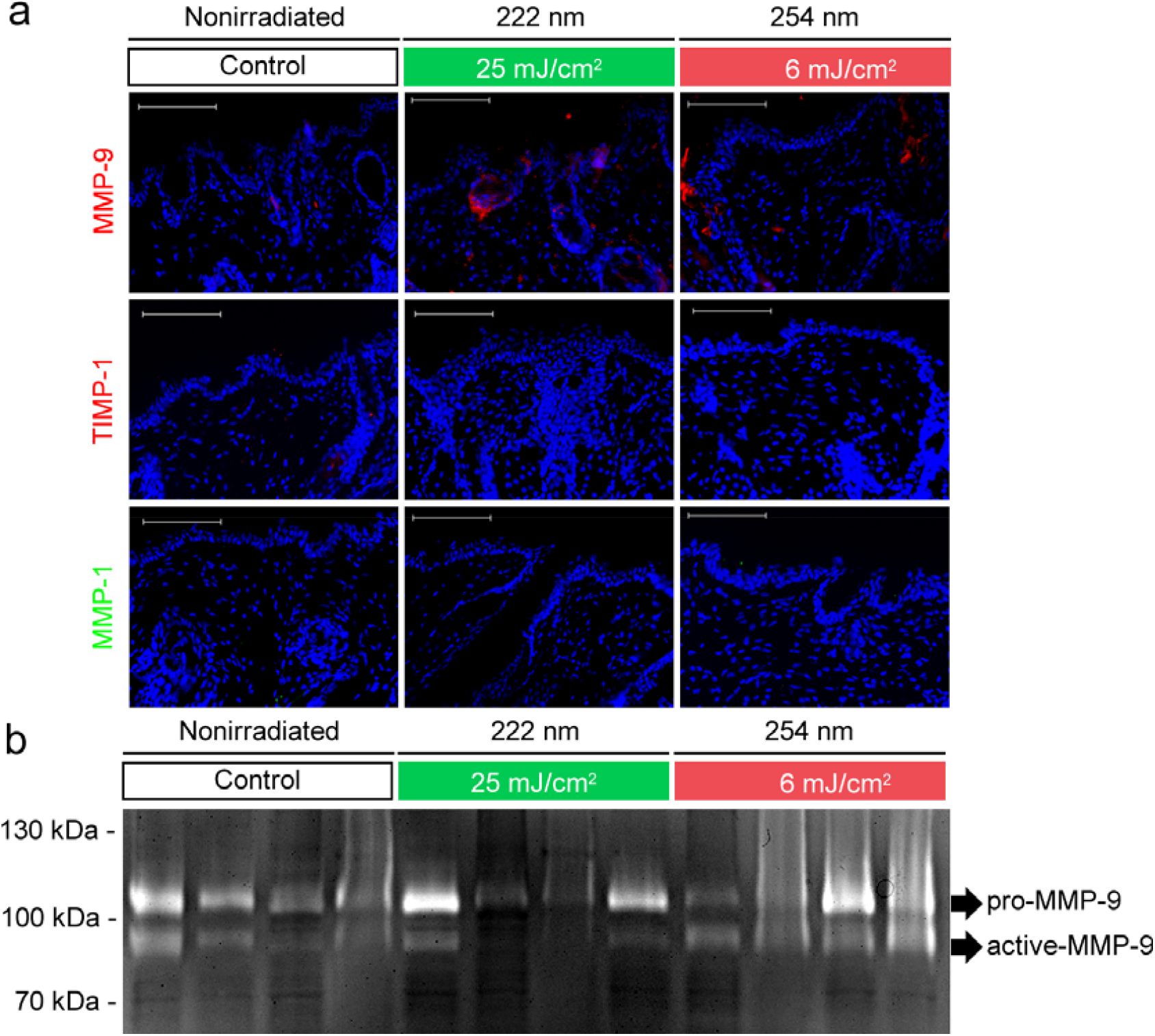
Chronic UV 222 nm and UV 254 nm irradiation increased only MMP-9 levels in the skin of HRS/J mice. HRS/J mice were nonirradiated or irradiated with 25 mJ/cm2 UV 222 nm or 6 mJ/cm2 UV 254 nm for eight hours, five days a week, for a total of 40 days. (a) Anti-MMP-1, anti-MMP-9, and anti-TIMP-1 staining were evaluated by immunofluorescence (anti-MMP-1 in green and anti-MMP-9 and anti-TIMP-1 in red; nuclei were stained with DAPI, in blue). Images are representative of the treatment group. (b) Zymography performed with a gelatin gel revealed increased levels of pro-MMP-9 and active MMP-9 after irradiation with 6 mJ/cm2 UV 254 nm. Pro-MMP-9 was detected at approximately 100 kDa, and active-MMP-9 was detected at approximately 90 kDa. Arrows indicate pro and active versions of MMP-9.

### 3.4 UV 222 nm and UV 254 nm increase ROS formation in the skin

As indicated in our proteomics study and the literature, UV-C radiation can generate oxidative stress in cells [1]. ROS can also increase the activity of MMPs [43], with both processes related to skin aging. To verify whether UV 222 nm and UV 254 nm could generate ROS in the RHS, we used the fluorescent probe DCFH_2_-DA, which, upon conversion to DCF inside cells, emits fluorescence at 488 nm. The RHS was irradiated with 6 mJ/cm², 25 mJ/cm², and 1500 mJ/cm² UV 254 nm or UV 222 nm and then immediately frozen. Skin irradiated with 1500 mJ/cm² UV 254 nm generated a stronger ROS signal than skin irradiated with the same dose of UV 222 nm (Figure 6a-b), corroborating our proteomic findings (Figure 3c). On the other hand, skin irradiated with 25 mJ/cm² UV 222 nm presented higher ROS levels than skin irradiated with 6 mJ/cm^2^ UV 254 nm (p-value < 0.0001, Tukey test) (Figure 6a-b). In summary, UV 222 nm generated a significantly higher ROS signal in irradiated skin than UV 254 nm at the respective TLVs.

**Fig. 6.**
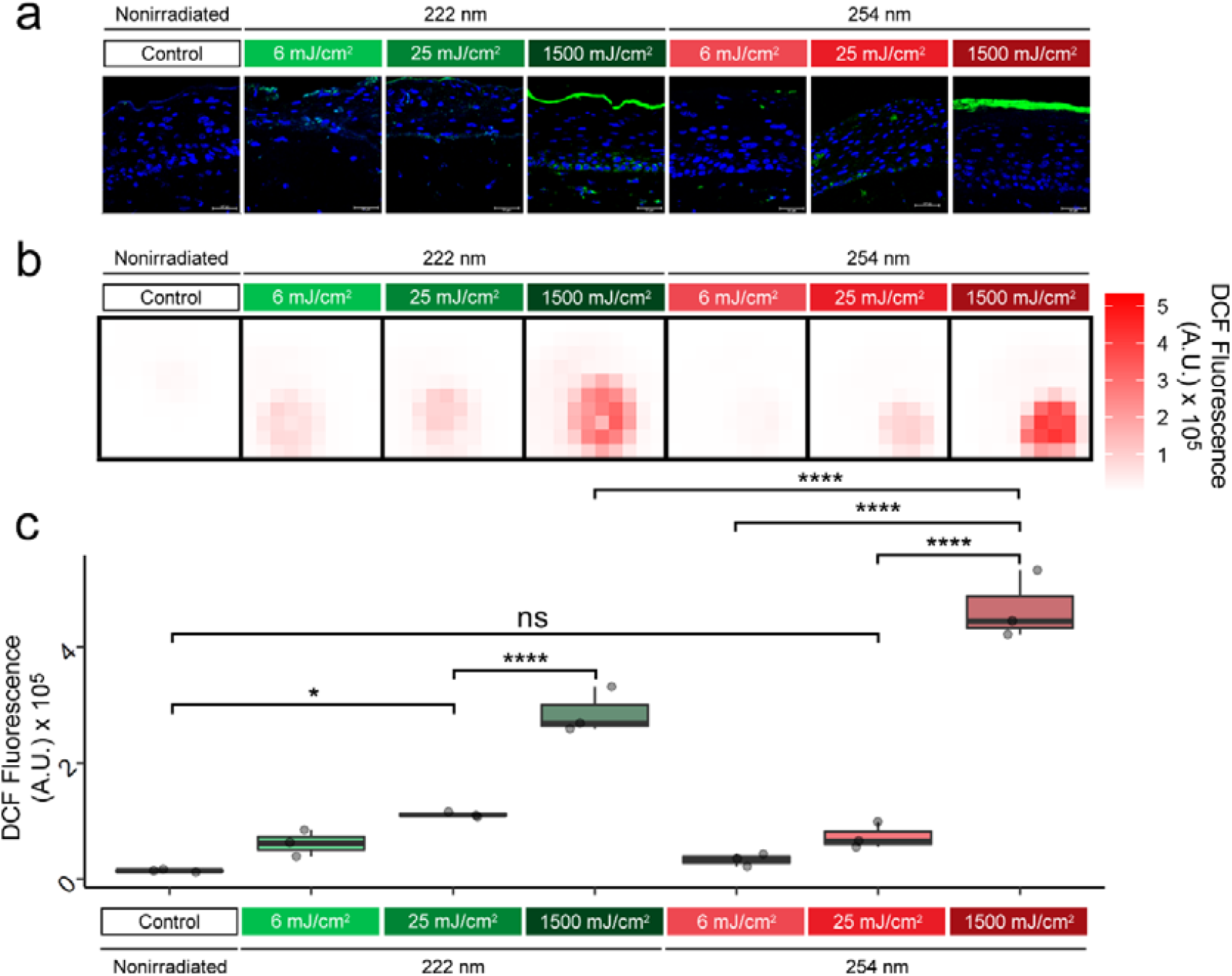
UV 222 nm and 254 nm increased ROS levels in the RHS. (a) Cryosections of RHS were irradiated with either 6, 25, or 1500 mJ/cm^2^ doses of either UV 222 nm or UV 254 nm (TLVs) and immediately incubated with a DCFH2-DA intracellular probe. Inside the cells, the probe is converted to DCF, which emits fluorescence at 488 nm (green). Images are representative of the treatment group. (b) The heatmap representation of the total fluorescence intensity of each skin area was obtained using a plate reader adapted to 24-well plates. (c) The maximum intensity of DCF fluorescence per well for biological replicates. P-values were derived from ANOVA followed by Tukey’s honestly significant difference test, with comparisons of different doses of each type of irradiation (UV 222 nm or 254 nm) at the maximal doses for both wavelengths.

## 4 Discussion

Far-UV-C radiation (200-230 nm) is a region of the UV-C spectrum not traditionally used for disinfection, although it has been reported to be an effective antimicrobial and antiviral agent. Notably, whereas exposure to conventional UV-C (250-280 nm) at germicidal doses is potentially hazardous, biophysical and experimental evidence suggests this is not the case for far-UV-C since greater absorption by protective surface layers results in much less damage to the skin and eye while maintaining disinfection efficacy.

Several recent papers have focused on krypton-chloride excimer (KrCl) lamps that exhibit a dominant peak at ∼222 nm [12,16,24]. Many studies have evaluated the impact of UV 222 nm on human and animal cells, skin, and eyes [18] and parameters such as cell number/viability, epidermal thickening, CPD and 6,4-PP levels, and erythema formation. Most of the presented studies concluded that human and animal cells could tolerate far-UV-C doses higher than 150 mJ/cm² without the abovementioned cell and tissue damage [18]. This irradiation dose is higher than the dose necessary to inactivate several microorganisms by several orders of magnitude [18]. Therefore, preventing surgical site infections with far-UV-C irradiation [12] seems plausible and attractive.

Despite the positive results reported in the available literature, the safety of far-UV-C irradiation has not yet been proven conclusively. In this work, we evaluated for the first time the effect of UV 222 nm on the proteome of the irradiated epidermis and consequent alterations in cellular pathways, ROS generation, and metalloproteinase levels and activity. For the first time, we also performed a 40-day *in vivo* study of hairless mice at the allowed UV 222 nm occupational safety dose (25 mJ/cm² per eight hours), with UV 254 nm as a control.

We confirmed, as previously published [12,16,20,21], that UV 254 nm generates more and deeper cell layer DNA lesions than UV 222 nm in both *in vitro* RHS models and HRS/J mouse skin after acute exposure at the highest tested dose (1500 mJ/cm²). Furthermore, while the DNA lesions generated by UV 222 nm slough off more quickly after 48 h of exposure, skin irradiated with UV 254 nm still presents DNA lesions and important morphological alterations after 48 h, indicative of more severe skin damage.

Our microscopy, proteomic, MMP, and ROS detection studies with the RHS revealed that, as expected, UV 254 nm exposure led to visible alterations in irradiated skin, with apparent disorganization of the dermis and epidermis, the potential to generate an inflammatory response, and extracellular matrix degradation, with an increase in MMP activity, which may all be related to increased levels of ROS. Skin exposed to UV 222 nm showed a proteomic profile closer to that of nonirradiated skin than to that of UV 254 nm-irradiated skin in terms of oxidative stress (although we could directly detect ROS after UV 222 nm irradiation even at the TLV dose), and the inflammatory response. In this sense, it is possible that although UV 222 nm can promote ROS generation and increase MMP levels, counteracting processes, such as TIMP-1 activity, may be enough to preserve the tissue. Of note, integrins, fibril formation, wound healing, and collagen formation processes had lower scores in RHS 48 h post-irradiation with UV 222 nm than in nonirradiated or UV 254 nm-irradiated skin. Finally, although the alterations in the abovementioned pathways caused by UV 222 nm may, in the short term, protect skin from acute degradation following high UV 222 nm doses, they raise some concerns about the ability of long-term and direct exposure to alter the capacity of the skin to regenerate. Such impairment may lead to premature aging, but this assumption requires further study.

## 5 Conclusions

Our study evaluated for the first time the proteome and cellular pathway alterations induced by UV 222 nm irradiation in an artificial skin model, and we showed that this type of radiation minimally alters processes related to reactive oxygen species and the inflammatory response compared to the more frequently used germicidal UV 254 nm lamp. For the first time, we also performed a 40-day *in vivo* exposure assay on hairless mice using the occupational threshold value of UV 222 nm and showed that UV 222 nm caused minor damage to the skin of exposed mice. However, alterations of pathways related to skin regeneration raise concerns about the possibility of premature aging after long-term and direct exposure to UV 222 nm.

## 6 Author contributions

RSNT: Conceptualization; Data curation; Formal analysis; Investigation; Methodology; Validation; Visualization; Roles/Writing - original draft; Writing - review & editing.

DA: Software; Data curation; Formal analysis; Methodology; Roles/Writing - original draft; Writing - review & editing.

AG: Methodology; Roles/Writing - original draft; Writing - review & editing.

ENL: Methodology; Writing - review & editing.

ASJ: Methodology; Writing - review & editing.

RD: Methodology; Software; Data curation; Writing - review & editing.

ACC: Software; Data curation.

REM: Funding acquisition; Resources; Writing - review & editing.

MC: Software; Data curation.

ALBA: Methodology; Writing - review & editing

AFPL: Resources; Methodology; Software; Data curation; Writing - review & editing

SMDG: Conceptualization; Visualization; Funding acquisition; Project administration; Resources; Supervision; Roles/Writing - original draft; Writing - review & editing.

## 7 Acknowledgments

This research used the facilities Brazilian Biosciences National Laboratory (LNBio), both part of the Brazilian Centre for Research in Energy and Materials (CNPEM), a private non-profit organization under the supervision of the Brazilian Ministry for Science, Technology, and Innovations (MCTI). The Mass Spectrometry Laboratory staff are acknowledged for their assistance during the experiments (Proposal number MAS-20220040). This work was supported by Empresa Brasileira de Pesquisa e Inovação Industrial (EMBRAPII), Instituto Tecnológico da Vale (IT-Vale), Vale S/A [grant number PCNP-2007.0015], and FAPESP (2018/16453-8). We acknowledge CIBFar (FAPESP CEPID 2013/07600-3) for full support. We acknowledge the researchers Verônica Teixeira and Wanderley Pedroso da Graça, for the device development and characterization. Professors Dr. Lorena Rigo Gaspar Cordeiro and Dr. Silvya Stuchi Maria-Engler for the artificial skin model technology.

## 8 Data availability

The mass spectrometry proteomic data have been deposited in the ProteomeXchange Consortium (http://proteomecentral.proteomexchange.org) via the PRIDE partner repository [44] with the dataset identifier PXD037726.

## 9 Code availability

R code used to prepare protemic data is available at GitHub repository (https://github.com/douglasadamoski/farUVC).

