## Supplemental Figures for "Different biological effects of exposure to far-UVC (222 nm) and near-UVC (254 nm) irradiation"

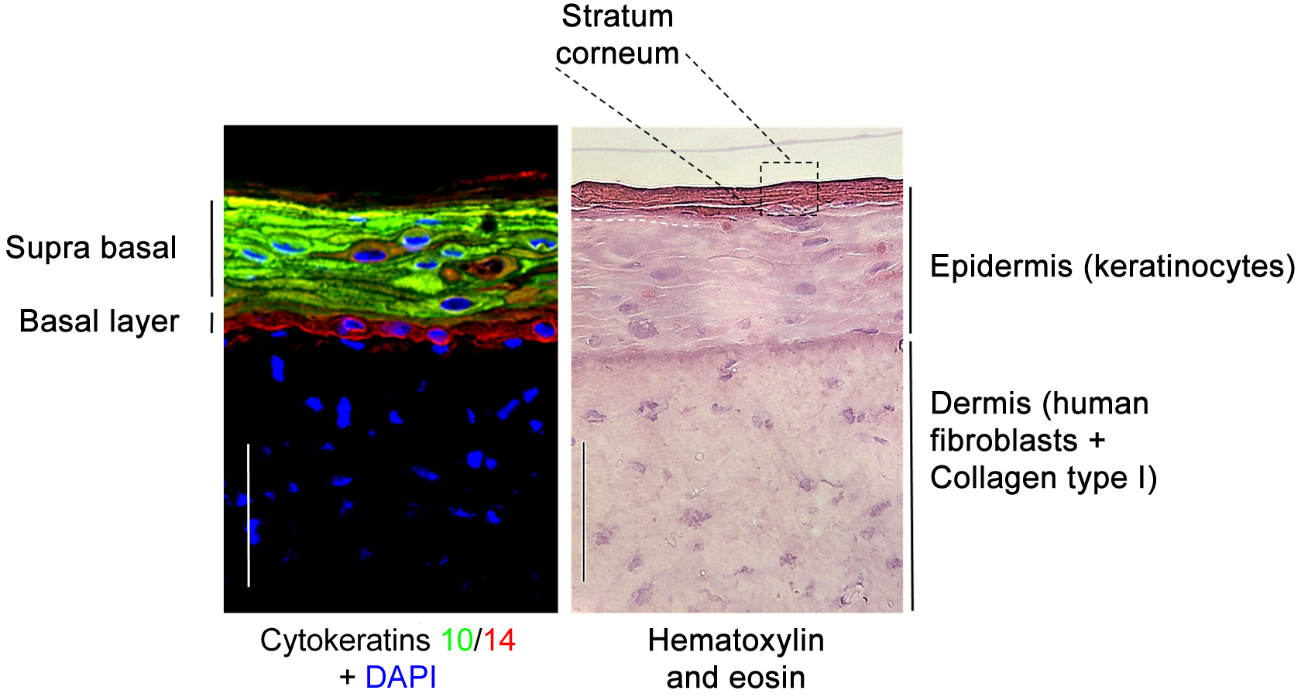

**Supplementary Fig. 1. Histological characterization of the RHS.** On the left, immunofluorescence showing cytokeratin 10 (green), which is mainly found in more differentiated keratinocytes in the suprabasal layer, and cytokeratin 14 (red), which is mainly found only in the proliferative cells of the basal layer; nuclei were stained with DAPI (blue). On the right, hematoxylin shows fibroblasts well distributed in the dermis and keratinocytes in the stratified epidermis. Eosin stained the stroma and the stratum corneum (containing cells without nuclei). Images are representative of RHS.

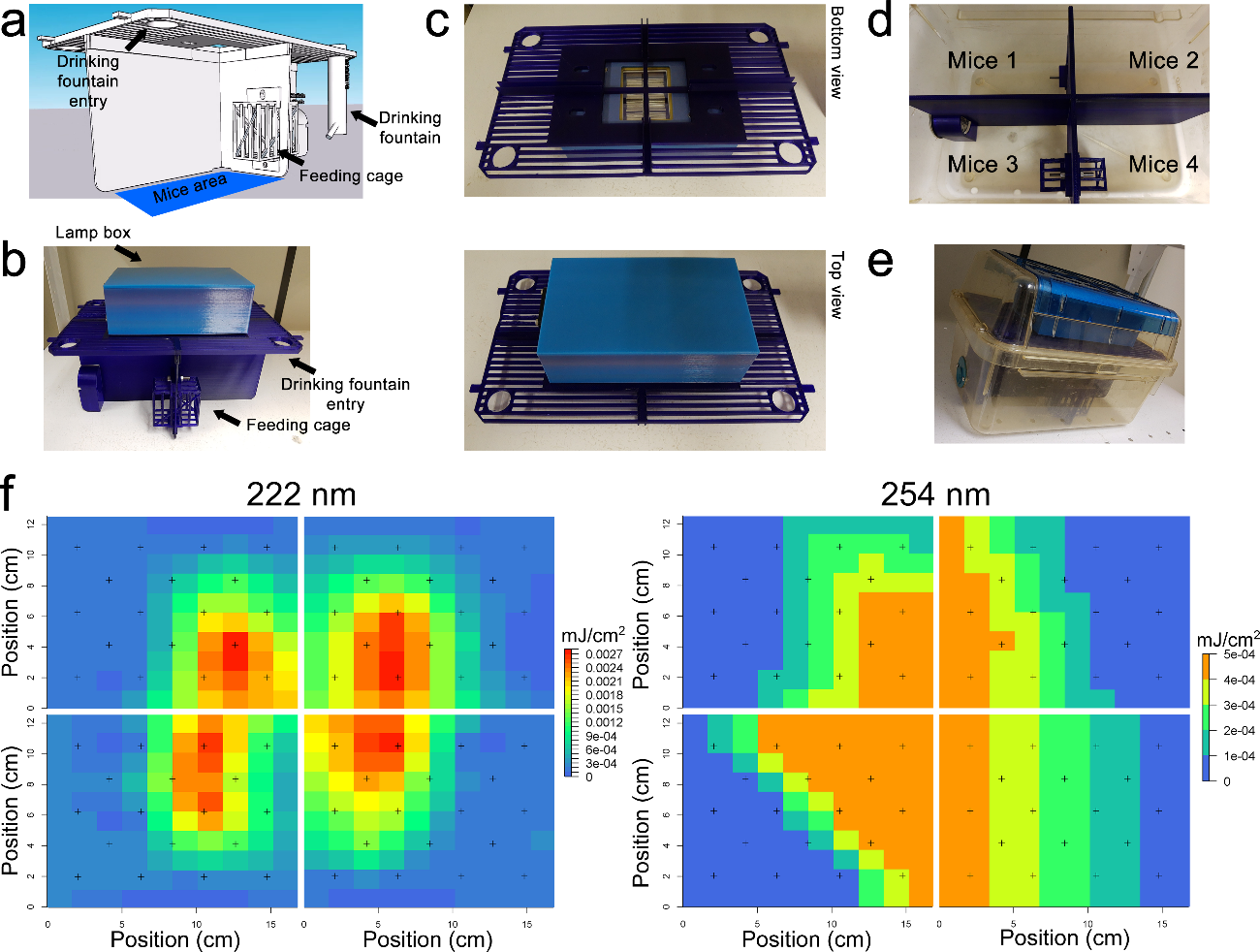

**Supplementary Fig. 2. In-house constructed *in vivo* chronic irradiation device.** (a) The 3D projection of the compartments shows in detail the cage with independent food and water dispensers. (b) The device was 3D printed by extrusion using polylactic acid filament in an S3X Printer (Sethi3D) with printed lamp support (light blue box on the top); (c) bottom and top detailed view of the ventilation grid fin with UV 222 nm lamp holder; (d) Printed divider placed inside of the animal cage (without the lid) (e). Fully assembled animal microisolator. (f) Heatmap of the UV 222 nm (on the left) and UV 254 nm (on the right) radiance distribution inside each animal compartment.

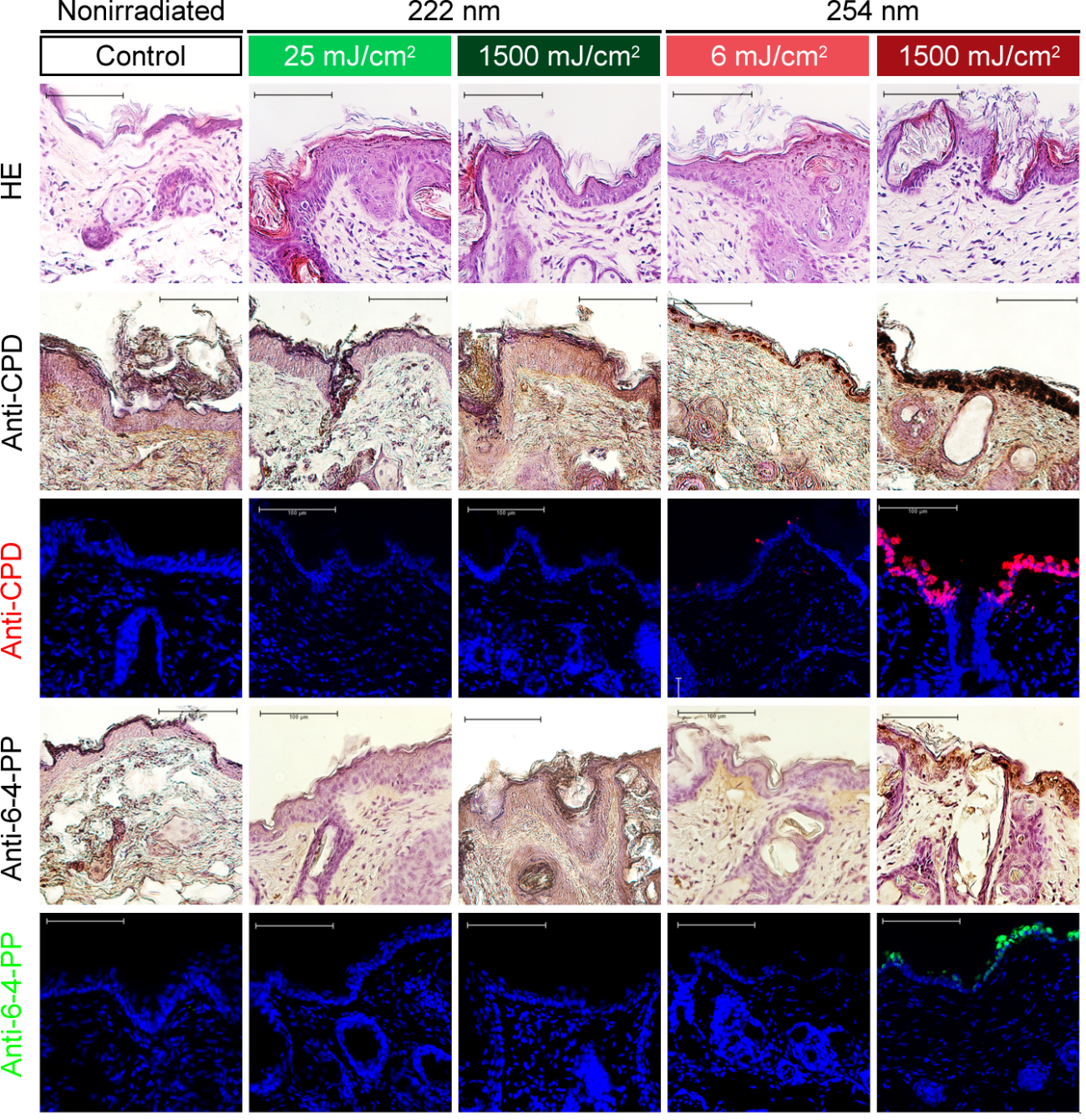

**Supplementary Fig. 3. Morphological changes and dimer formation in the skin of HRS/J mice after acute exposure.** HRS/J mice were nonirradiated (control, white box) or irradiated with 6, 25, 500, 1500 mJ/cm^2^ UV 222 nm (green box) or UV 254 nm (red box). HE: hematoxylin and eosin staining; anti-CPD and anti-6-4-PP staining by IHC and IF techniques. Images are representative of the treatment group. Bar size, 100 µm.

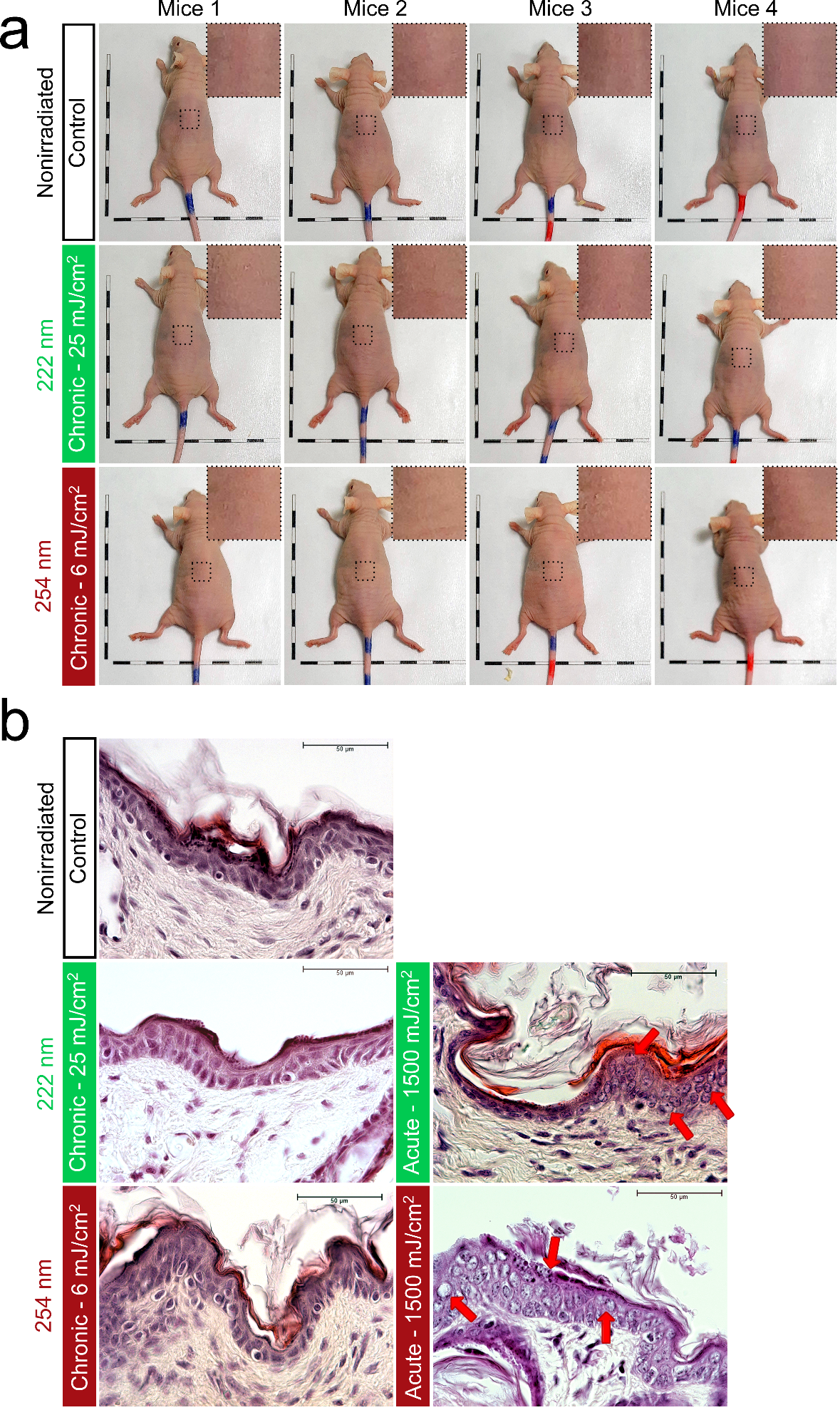

**Supplementary Fig. 4. HRS/J mice do only show histological effects of short- and long-term exposure to UV 222 nm.** HRS/J mice were nonirradiated (control, white box) or irradiated with UV 222 nm (green boxes) or UV 254 nm (red boxes). (a) Macroscopic view of mice subjected to chronic exposure after euthanasia. Black boxes represent 1 cm. Dashed boxes represent a magnified area in the upper right corner of each mouse. (b) Hematoxylin and eosin staining of skin from both chronic and acute exposures. Red arrows indicate abnormal hydropic degeneration focus cells in 1500 mJ/cm^2^ from both lamps. Bar size, 50 µm.

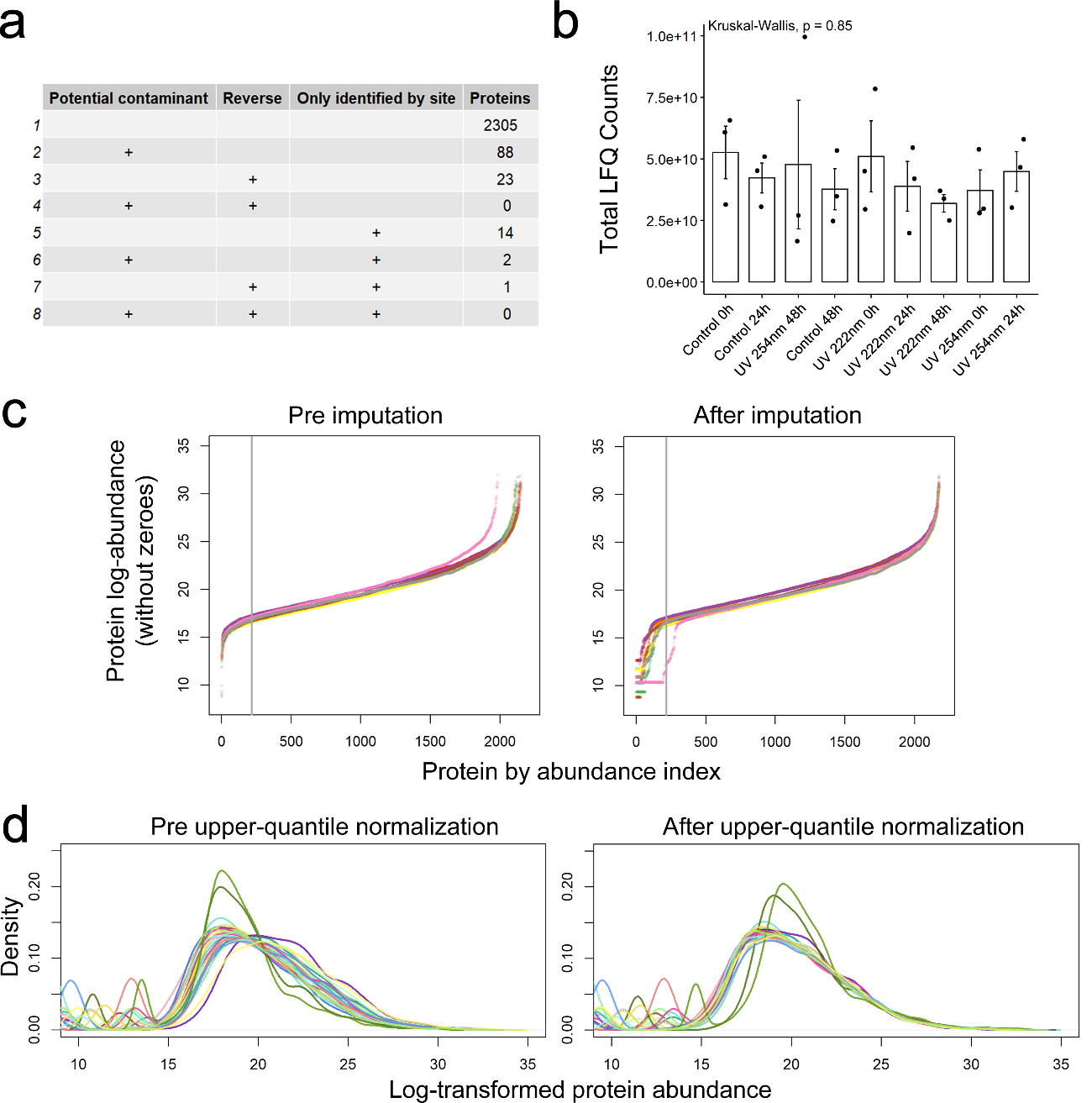

**Supplementary Fig. 5. Label-free proteomics analysis of the UV-exposed RHS, including quality control, missing data imputation, and normalization.** (a) MaxQuant category quality filtering. (b) Sum of label-free quantification (LFQ) counts for all thresholded proteins under all conditions. Error bars represent the standard deviation. Each dot is one sample. The Kruskal‒Wallis variance test was applied. (c) Distribution of protein abundances (as Log_2_-LFQ) ordered by abundance index before (left) and after (right) imputation. (d) Distribution of the imputed log_2_-LFQ before (left) and after (right) upper-quantile normalization.

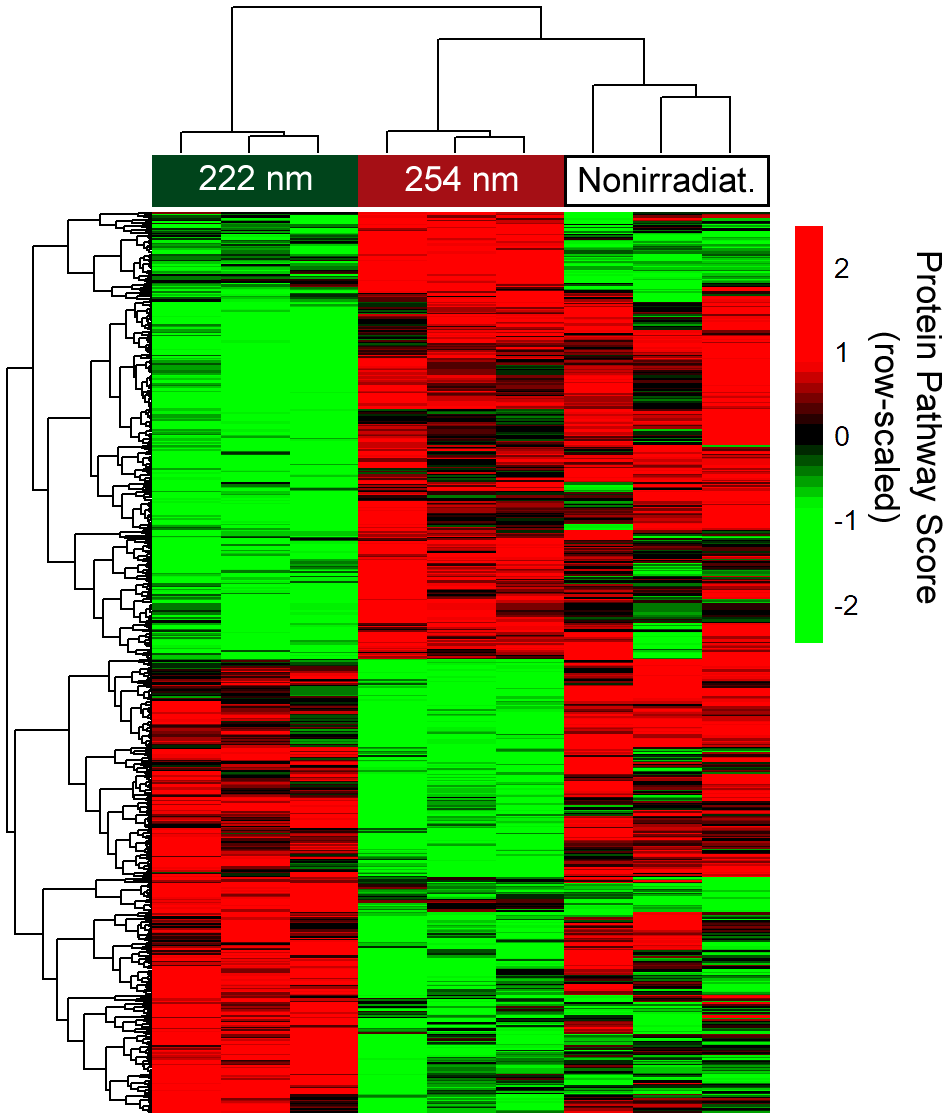

**Supplementary Fig. 6. Proteomics analysis after UV 222 nm and 254 nm exposure in an RHS model shows multiple altered pathways compared to the control.** Heatmap of 635 single-sample protein set scores from MSigDB showed statistically significant differences (p < 0.05, t-test) after 48 h of exposure in the UV 222 nm and 254 nm comparison. Rows and columns were clustered using the complete method, and distances were calculated using Pearson correlation.

**Supplementary Table.** Raw and processed data from proteomics studies.
